# Breeding Program Optimization for Genomic Selection in Winter Wheat

**DOI:** 10.1101/2020.10.07.330415

**Authors:** Megan Calvert, Byron Evers, Xu Wang, Allan Fritz, Jesse Poland

## Abstract

Developing methodologies in the fields of phenomics and genomic prediction have the potential to increase the production of crop species by accelerating germplasm improvement. The integration of these technologies into germplasm improvement and breeding programs requires evidence that there will be a direct economic benefit to the program. We determined a basic set of parameters, such as prediction accuracy greater than 0.3, the ability to genotype over 7 lines for the cost of one phenotypic evaluation, and heritability levels below 0.4, at which the use of genomic selection would be of economic benefit in terms of genetic gain and operational costs to the Kansas State University (KSU) winter wheat breeding program. The breeding program was then examined to determine whether the parameters benefitting genomic selection were observed or achievable in a practical sense. Our results show that the KSU winter wheat breeding program is at a decision point with regards to their primary means of selection. A few operational changes to increase prediction accuracy would place the program in the parameter space where genomic selection is expected to outpace the current phenotypic selection methodology at a parity of the operation cost and would be of greatest benefit to the program.

## Introduction

Technologies are constantly evolving and growing in today’s ever-changing world. The introduction of new technologies into an essential service and the development of new crop varieties can be challenging and costly (Moose and Mumm 2008). New technologies must be carefully evaluated to determine if they provide a significant enough advantage to adjust proven current practice. There are many technologies that appeared to have great promise, such as quantitative trait loci (QTL), and marker-assisted selection (MAS), and yet were not beneficial in a practical sense to breeding programs at the time (Bernardo 2008). Yet with a growing global population, erratic environmental conditions, and a finite-amount of arable land, the introduction of new technologies into our crop development systems is essential. The determination of which technologies to implement and the most efficient way to do so is a constant challenge facing plant breeders.

Wheat (*Triticum aestivum*) is one of the top three field crops planted in the USA, behind soybeans and corn, with 1.9 billion bushels produced in 2019 (Bond 2020). Wheat acres and production in the USA have been in a decline since the 1980s as a result of international competition, changing economic conditions and production practices. This is despite the increasing demand for wheat due to growing global populations, which is only expected to increase over the coming century. It is estimated that a 2% increase in yearly grain yields are required to meet these demands (Bassi et al. 2016). Currently there are no reports of wheat breeding programs achieving this level of gain, making wheat an important candidate for breeding technologies designed to accelerate variety development.

Genomic prediction (GP) is one of the new technologies that is showing a great deal of promise to assist in crop-variety production. GP involves the use of genome-wide markers to predict the breeding value of individuals in a population (Meuwissen, Hayes, and Goddard 2001). This is done by genotyping and phenotyping a training population which is used to establish a model for the trait of interest. This model is then used to calculate genomic estimated breeding values (GEBVs) of a population for which only genotype information is available. GP is already a common technique used in animal breeding due to the benefit of being able to predict a phenotype without having to observe the phenotype, for example the milk yield of a bull’s offspring (Hayes et al. 2009; Georges, Charlier, and Hayes 2019).

Plant breeding programs have yet to fully utilize GP for a number of reasons. Breeding programs have to test the same experimental line in multiple locations due to the need to select varieties that are stable across environments. This negates some of the benefit that GP supplies to animal breeding, in which the same genotype cannot be tested under multiple conditions. The genotype x environment (GxE) variation is often high in plant breeding populations due in part to the large weather differences experienced between locations. This significantly decreases the accuracies of most GP models (Dawson et al. 2013; Heslot et al. 2012). Previous GP models did not account for GxE interactions which limited inference and selection decisions in plant breeding. Newer genomic prediction models can now take into account these GxE interactions and are showing greater accuracy in plant breeding situations (Burgueño et al. 2012; Pérez-Rodríguez et al. 2017; Montesinos-López et al. 2016).

The accuracy of GP models has shown some improvement when a covariate is included (Crain et al. 2018; Rutkoski et al. 2016). This has had some success when including high-throughput phenotyping (HTP) data that is taken throughout the course of the growing season. HTP techniques have the ability to measure thousands of phenotypes accurately in a short period of time. Many of the HTP techniques used in breeding programs take advantage of new developments in remote sensing, uncrewed aerial system (UAS), and sensor technology to measure reflectance and temperature phenotypes. These are all phenotypes that have been shown to be correlated with yield and as such could be effective secondary targets for high-throughput phenotyping and indirect selection (Elliott and Regan 1993; Blackmer et al. 1996; Curran et al. 1983).

In addition to the use of correlated traits, the efficiency of GP is also affected by the stage of the breeding program when it is implemented. Primarily, GP is used to predict the breeding value which is comprised of the additive genetic variation. The greatest advantages of GS will be found when the selection candidates still encompass the most additive genetic variation for the phenotype of interest (Bassi et al. 2016). This is more likely to occur in earlier stages of a breeding program as it is easily selected for, while later stages of a breeding have progressed through strong selection and are more likely to select for other epistatic genetic variation on the performance of the line *per se*.

Another factor in addition to stage of implementation that has hindered the adoption of GP in plant breeding programs are the costs and complex logistics associated with it. The development and establishment of genotyping practices in a plant breeding program requires an investment in infrastructure and training (Moose and Mumm 2008). Along with genetic marker costs, the establishment and maintenance of an adequate training population adds additional costs to the GP protocols. These costs may not be offset by the gains that the program could potentially achieve using GP (Bassi et al. 2016; Jarquín et al. 2017).

In this study we examined the implementation of GP in a wheat breeding program to specifically examine 1) what parameters/infrastructure is required to implement GP as a primary selection strategy, and 2) are those parameters being met in the KSU public breeding program?

## Method and Materials

### Cost-Benefit Simulation

A simulated breeding program was used to determine the expected genetic gain when basing selection of material exclusively on observed line performance *per se*, or exclusively on prediction of breeding values. The simulation assumes that a breeding program does not use a combination of phenotypic selection and genomic selection at the same time. In consultation with the Kansas State Hard Winter Wheat Breeding Program, referred to as the KSU breeding program from here on, an appropriate range of cost estimates were determined, as well as sizes of the program as determined by the number of lines developed and evaluated (Supplementary Table 1). For simplicity looking at a single stage of selection in the program, the simulated values assume that the cycle length for each selection scheme is the same and covers the period of the program before the Advanced Yield Nursery (AYN) stage. It also assumes that the number of lines that will be selected for the AYN stage are the same regardless of which selection method is used.

The cost of line developments was estimated to be between $4-$30 per line which covers the initial cross and formation of an inbred line. This can either be done by several years of inbreeding or by the formation of double haploid lines. This was applied as a fixed cost needed to develop the initial population for selection regardless of which selection method was used.

### Phenotypic Selection

The costs of phenotypic observation plots are estimated to be between $12-$40 per plot and evaluated within this range at $5 increments. To obtain an accurate phenotypic measurement, replications of the experimental line need to be planted and phenotyped. Depending on the structure of the program this may mean several replications at a single site, or fewer replications at several sites. The maximum number of experimental breeding lines was calculated as:

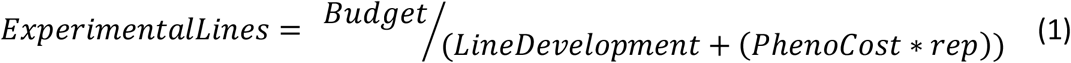

where *ExperimentalLines* is the number of lines that would be advanced to the phenotyping stage of selection, the *Budget* is the total monetary budget for that stage of the breeding program, *LineDevelopment* is the cost of advancing a single cross to an inbred experimental line for evaluation, *PhenoCost* is the total cost to obtain the phenotype of interest including labor and other miscellaneous operation costs, and *rep* is the number of replications of each experimental line that are planted in field and will require phenotyping.

The number of experimental breeding lines was used as the population size when estimating other population parameters.

### Genomic Selection

The cost of genotyping a single line for prediction was estimated to be between $1-$10 and evaluated within this range at $1 increments. The maximum number of experimental lines that can be genotyped for prediction was calculated as:

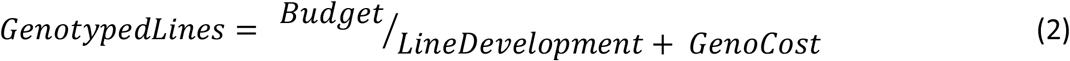

Where *GenotypedLines* is the number of lines that would be evaluated by genotyping, the *Budget* is the total monetary budget for that stage of the breeding program, *LineDevelopment* is the cost of advancing a single cross to an inbred experimental line for evaluation, and *GenoCost* which is the cost of genotyping a single line including labor and other operational costs. It is assumed that the genotyping is only performed once, and that replication is not required.

The number of possible genotyped lines was then used as the population size when estimating other population parameters.

### Simulation Details

The selection methods were compared for every possible combination of number of experimental lines phenotyped and number of experimental lines genotyped, based on the ratio between the correlated response to selection, and the response to selection (Falconer and Mackay 2009).

The expected response to selection for the primary trait was calculated by:

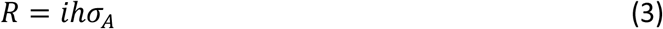

Where *i* is the intensity of selection, *h* is the square-root if the narrow-sense heritability of the primary trait, and *σ*_*A*_ is the standard deviation of the additive genetic variance (Falconer and Mackay 2009).

For this study, the GEBVs are assumed to be the secondary trait that is correlated to the primary trait, which would be through experimental observation plots. The correlated response to selection is calculated by:

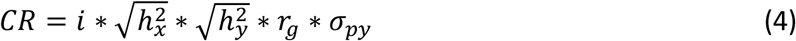

Where *i* is the intensity of selection, 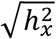 is the square-root of the narrow-sense heritability of the response trait, 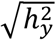 is the square-root of the narrow-sense heritability of the secondary trait, *r*_*g*_ is the additive genetic correlation between the traits, and *σ*_*py*_ is the standard deviation of the phenotypic variation for the secondary trait (Falconer and Mackay 2009).

A comparison between the indirect response to selection and the expected response to selection is best demonstrated as (Falconer and Mackay 2009):

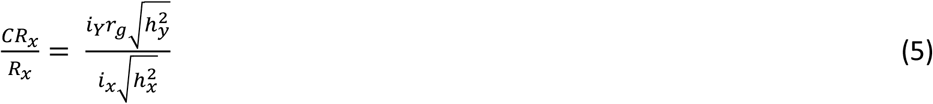

Where *i*_*Y*_ is the selection intensity on the secondary trait, *r*_*g*_ is the additive genetic correlation between the primary and secondary trait, 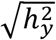 is the square-root of the narrow-sense heritability of the secondary trait, *i*_*x*_ is the selection intensity on the direct trait, and 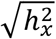 is the square-root of the narrow-sense heritability of the response trait.

It was assumed that the individual phenotypes were made up of a genetic portion and an environmental portion. The distribution of each of these portions was assumed to be a random normal with a mean of 0, and a standard deviation of 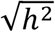 or 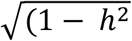 for the genotypic and environmental proportions, respectively. A sample the size of the number of lines that were possibly genotyped was taken from this distribution for each individual to give the overall phenotypic distribution. This overall phenotypic distribution was used to determine the intensity of selection (*i*) in terms of the standard deviation and the selection differential with the msm package (Jackson 2011).

The narrow sense heritabilities for the response trait and the additive genetic correlation were set, with testing the ranges of 0 and 1 at increments of 0.1 each. The narrow-sense heritability of the secondary trait, the genotyping, was assumed to be 0.95. This is under the assumption that the genotypes are inherited almost exactly as they are sequenced and that there are only a few genotyping errors. The ratio between the correlated response and the expected response to selection was plotted against the ratio between number of experimental lines phenotyped and number of lines genotyped.

### Plant Material

The KSU Breeding Program breeds hard red winter wheat for a large area which contains different mega-environmental conditions. A subset of 5 locations within the same Kansas mega-environment based on breeder knowledge, Belleville (BEL), Gypsum (GYP), McPherson (MP), Hutchinson (HUTCH) and Manhattan (MANH), were selected for analysis in Kansas between 2016 and 2019. This resulted in 1989 experimental lines being examined over the course of 4 years. These sites contain trials from the Preliminary Yield Trials (PYN, primary F_5:7_) and the advance yield nursery (AYN, primarily F_5:8_). The planting, harvest dates and trial size are provided in Table 1. These locations, excluding Manhattan and Hutchinson, are located in farmers’ fields under typical grower management practices. The PYN and AYN trials were all planted in six-row plots of 1.5m by 4.5m. The PYN are planted in a modified augmented design with one replicate of the experimental line per location (Federer and Raghavarao 1975). Plant checks are planted across whole rows and columns in the trial, and sub-block checks are assigned randomly within each block. The AYN is made up of lines selected from the PYN trials. The lines are planted using two replicated α-lattice designs (Patterson and Williams 1976). All 5 locations were planted each year but if a site experienced extreme environmental variation from the normal climate it was not harvested, providing an unbalanced set of data.

**Table 0.1.** **Summary of trials planted between the 2016 and 2019 seasons across locations. A summary is given by season, trial and location with the number of individual plots, experimental lines, date of planting, date of harvest, the mean grain yield in t/ha and the standard deviation of the grain yield**.

| Season | Trial | Location | Plots | Lines | Planted | Harvested | Mean | SD |
| --- | --- | --- | --- | --- | --- | --- | --- | --- |
| 2015/2016 | AYN | MP | 40 | 75 |  |  | 5.050 | 0.9 |
|  |  | MANH | 240 | 75 |  |  | 3.206 | 0.651 |
|  |  | HUTCH | 320 | 109 |  |  | 4.139 | 0.92 |
|  |  | GYP | 40 | 109 |  |  | 5.739 | 0.726 |
|  | PYN | MP | 504 | 439 |  |  | 5.113 | 1.21 |
|  |  | MANH | 504 | 379 |  |  | 3.720 | 0.657 |
|  |  | HUTCH | 594 | 516 |  |  | 3.770 | 0.886 |
|  |  | GYP | 593 | 506 |  |  | 5.421 | 0.702 |
| 2016/2017 | AYN | MP | 315 | 91 | 10/11/16 | 20/06/17 | 5.933 | 1.00 |
|  |  | MANH | 320 | 91 | 10/18/16 | 22/06/17 | 5.822 | 0.768 |
|  |  | HUTCH | 168 | 97 | 10/12/16 | 21/06/17 | 6.242 | 0.845 |
|  |  | BEL | 48 | 36 | 10/19/16 | 28/06/17 | 5.110 | 0.776 |
|  | PYN | MP | 126 | 120 | 10/11/16 | 20/06/17 | 4.805 | 1.015 |
|  |  | HUTCH | 108 | 102 | 10/12/16 | 21/06/17 | 6.319 | 0.925 |
|  |  | BEL | 8 | 8 | 10/19/16 | 28/06/17 | 4.5 | 0.385 |
| 2017/2018 | AYN | MP | 199 | 90 | 10/19/17 | 29/06/18 | 3.203 | 0.412 |
|  |  | MANH | 200 | 90 | 10/20/17 | 23/06/18 | 2.644 | 0.402 |
|  |  | HUTCH | 280 | 125 | 10/18/17 | 28/06/18 | 3.345 | 0.489 |
|  |  | BEL | 280 | 125 | 10/17/17 | 30/06/18 | 2.561 | 0.430 |
|  |  | GYP | 279 | 125 | 10/18/17 | 18/06/18 | 2.343 | 0.318 |
|  | PYN | MP | 323 | 282 | 10/19/17 | 29/06/18 | 3.048 | 0.490 |
|  |  | MANH | 414 | 366 | 10/20/17 | 23/06/18 | 2.880 | 0.431 |
|  |  | HUTCH | 414 | 366 | 10/18/17 | 28/06/18 | 3.236 | 0.435 |
|  |  | BEL | 287 | 251 | 10/17/17 | 30/06/18 | 2.198 | 0.412 |
|  |  | GYP | 286 | 250 | 10/18/17 | 18/06/18 | 1.973 | 0.278 |
| 2018/2019 | AYN | MANH | 80 | 40 | 11/01/18 |  | 5.798 | 0.535 |
|  |  | HUTCH | 212 | 99 | 10/24/18 | 01/07/19 | 4.925 | 0.563 |
|  |  | BEL | 130 | 63 | 10/24/18 | 17/07/19 | 4.965 | 0.897 |
|  | PYN | HUTCH | 504 | 454 | 10/24/18 | 01/07/19 | 4.623 | 0.794 |

### Phenotyping

Phenotypic information was collected either by combine for grain yield (GRYLD), by UAS for vegetation indices (VIs), or by hand using the Field Book application (Rife and Poland 2014) for plant height (PTHT). A DJI Matrice100 (DJI, USA) quadcopter UAS was equipped with a 5-band multi-spectral RedEdge camera (MicaSense Inc. USA) to collect plot-level reflectance values for each year, based on the standard protocols developed by The Wheat Genetics Lab at Kansas State University as laid out in Wang et al. (2018). The UAS data was collected throughout the course of the growing season, once every 7-10 days depending on weather conditions. Plant height was collected manually during the grain ripening stage before harvest. VIs were calculated from the reflectance values based on the protocols laid out in Wang et al. (2018) and given in Table 2.

**Table 0.2.** **Summary of vegetation indices used in the KSU breeding program across 5 years. For each index the formula and reference are provided**.

| **Abbr.** | **Index** | **Equation** | **Reference** |
| --- | --- | --- | --- |
| **GNDVI** | Green Normalized Difference Vegetation Index | $GNDVI = \frac{NIR - Green}{NIR + Green}$ | (Blackmer et al. 1996) |
| **NDRE** | Red Edge Normalized Difference Vegetation Index | $NDRE = \frac{NIR - RedEdge}{NIR + RedEdge}$ | (Elliott and Regan 1993) |
| **NDVI** | Normalized Difference Vegetation Index | $NDVI = \frac{NIR - Red}{NIR + Red}$ | (Curran et al. 1983) |

**Table 0.3.** **Genomic Prediction Accuracies as the correlation between BLUPs and GEBVs. The Pearson correlation coefficients is given with a 95% CI and the rank correlation between the GEBVs and the BLUPs**.

| **Training Population** | **Testing Population** | **Correlation** | **95% CI** | **Rank Correlation** |
| --- | --- | --- | --- | --- |
| Excluding 2015/2016 | 2015/2016 | 0.09 | [-0.004, 0.182] | 0.093 |
| Excluding 2015/2016 PYN | 2015/2016 PYN | 0.078 | [-0.026, 0.180] | 0.093 |
| Excluding 2015/2016 MP<br>MANH GYP | 2015/2016 MP<br>MANH GYP | 0.024 | [-0.08, 0.127] | 0.019 |
| Excluding 2016/2017 | 2016/2017 | -0.164 | [-0.336, 0.018] | -0.125 |
| Excluding 2016/2017 PYN | 2016/2017 PYN | -0.055 | [-0.249, 0.144] | 0.010 |
| Excluding 2016/2017 MP<br>MANH BEL | 2016/2017 MP<br>MANH BEL | -0.009 | [-0.206, 0.189] | 0.059 |
| Excluding 2017/2018 | 2017/2018 | 0.048 | [-0.049, 0.145] | 0.030 |
| Excluding 2017/2018 PYN | 2017/2018 PYN | 0.061 | [-0.046, 0.165] | 0.019 |
| Excluding 2017/2018 MP<br>MANH BEL | 2017/2018 MP<br>MANH BEL | 0.009 | [-0.097, 0.114] | -0.026 |
| Excluding 2018/2019 | 2018/2019 | -0.091 | [-0.186, 0.005] | -0.13 |
| Excluding 2018/2019 PYN | 2018/2019 PYN | -0.039 | [-0.135, 0.058] | -0.079 |
| Excluding 2018/2019<br>MANH BEL | 2018/2019<br>MANH BEL | 0.572 | [0.502, 0.634] | 0.537 |

### Genotyping

The 1989 lines in the study years were sequenced using genotype-by-sequencing on an Illumina Hi Seq2000 or Hi Seq2500 (Elshire et al. 2011). Single nucleotide polymorphisms (SNPs) were called with the Tassel software with the Chinese Spring wheat assembly v1.0 as a reference (Bradbury et al. 2007; Glaubitz et al. 2014; International Wheat Genome Sequencing Consortium et al. 2018). The final data set included 8182 SNPs that were selected for use passed one of three filtering criteria optimized for the wheat genome by Shrestha et al. (2020) that include Chi-square, Fisher’s test for independence, and the inbreeding coefficient, as well as having a minor allele frequency greater than 0.05 and missing less than 20% of the data. Missing SNPs were imputed with Beagle 5.1 (Browning, Zhou, and Browning 2018).

### Data Analysis

The Best Linear Unbiased Predictions (BLUPs) for each line were calculated for each individual year and for multiple years, including or excluding locations as needed. Where multiple years and locations are included, such as for GRYLD, the BLUPs were calculated by:

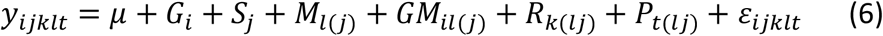

where *y*_*ijklt*_ is the phenotypic response variable,*μ* is the fixed overall mean, *G*_*i*_ is the random genotype effect for line i distributed as iid 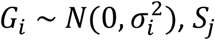 is the random effect for year *j* distributed as iid 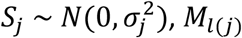 is the random effect for location *l* within year *j* distributed as iid 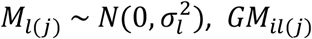 is the random genotype by location effect nested within year *j* distributed as iid 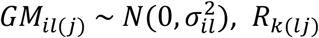 is the random effect of replication *k* within location and year distributed as iid 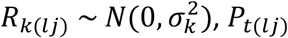 is the random fungal treatment effect *t* nested within year-location distributed as iid 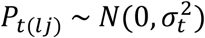, and *ε*_*ijklt*_ as the residual effect distributed as iid 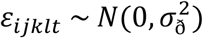.

When only a single year with multiple locations is included the BLUPs are calculated by:

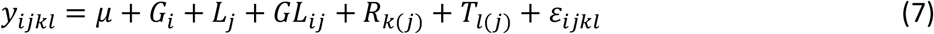

where *y*_*ijkl*_ is the phenotypic response variable,*μ* is the fixed overall mean,*G*_*i*_ is the random genotype effect for line i distributed as iid 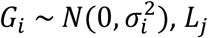 is the random effect of location *j* distributed as iid 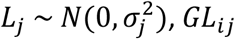 is the random effect of genotype by location distributed as iid 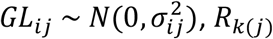 is the random effect of replication *k* nested in location *j* distributed as iid 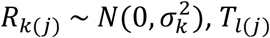 is the random effect of fungal treatment *l* nested within location *j* distributed as iid 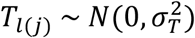,, and *ε*_*ijkl*_ is the residual effect distributed as iid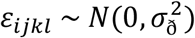.

The broad-sense heritability for all years and multiple year-locations was calculated as:

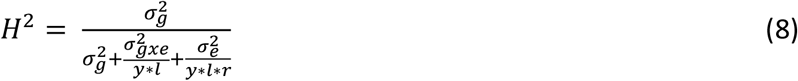

Where *σ*_*g*_ is the total genetic variance, *σ* _*gxe*_ is the variance contributed by the location and variety nested within year, *σ*_*e*_ is the residual environmental variance. As the data is unbalanced, y is the harmonic mean of the number of years planted, l is the harmonic mean of the number of locations planted within year, and r is the harmonic mean of the number of replications per location per year (Holland, Nyquist, and Cervantes-Martínez 2010).

The broad-sense heritability for individual year AYN and individual year-multiple location PYN was calculated by:

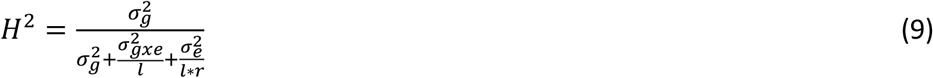

where 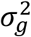 is the total genetic variance, 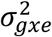 is the variance contributed by the genotype-location combination, and 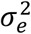 is the residual environmental variance. Similar to equation 8, *l* is the harmonic mean of the number of locations planted and r is the harmonic mean of the number of replications per location (Holland, Nyquist, and Cervantes-Martínez 2010).

The VI phenotypes were mainly considered on a year-location-trial basis. As such the BLUPs for the VI phenotypes for the AYN trials are calculated by:

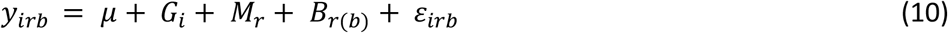

where *y*_*irb*_ is the phenotypic response variable, *μ* is the fixed overall mean,*G*_*i*_ is the random genotypic effect for line i distributed as iid 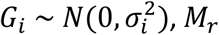 is the random effect of the replicate *r* distributed as iid 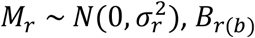, is the random effect of the experimental block nested within replicate distributed as iid 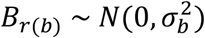, and *ε*_*irb*_ is the residual effect distributed as iid 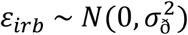. The broad-sense heritability was calculated for each year-location’s AYN that had VI phenotypes by:

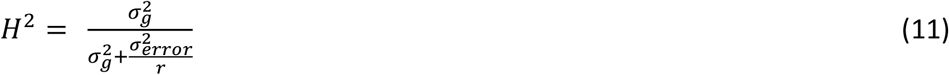

where 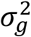 is the genetic variance, 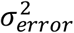 is the residual environmental variance and r is the number of replicates.

The BLUPs for the VI phenotypes for the PYN trials are calculated by:

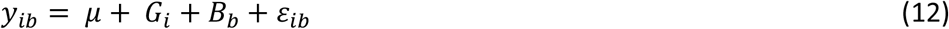

where *y*_*ib*_ is the phenotypic response variable, *μ* is the fixed overall mean, *G*_*i*_ is the random genotypic effect N(0,*σ*^2^), *B*_*b*_ is the random effect of the experimental block N(0,*σ*^2^), and *ε*_*ib*_ is the residual effect.

### Genome-wide Association Analysis

A principal component (PC) analysis of the genotypic information was conducted with the pcaMethods package (Stacklies et al. 2007). A genome-wide association analysis was performed for the VI BLUPs for each year-location-trial using the rrBLUP package (Endelman 2011). The kinship matrix was calculated using rrBLUP was included with 4 PCs to account for kinship and population structure respectively (Endelman and Jannink 2012). A Bonferroni correction was applied with α = 0.05 to determine significance.

### Genomic Prediction

Genomic prediction was performed with the rrBLUP package and the BGLR statistical package (Pérez and de los Campos 2014) in the R software environment. The rrBLUP package performs ridge regression (RR), and the BGLR package was used to perform a BayesC based prediction and a reproducing kernel Hilbert spaces regression (RKHS). A cross-validation strategy of 100 replications with 80% of the population in the training data set and 20% of the population in the predicted data set was used. The accuracy of the prediction was determined by the correlation between the predicted and observed values. Across year and location predictions were also performed where years, year-trial, and year-location combinations that were not included in the training population are predicted from the remaining data. An example of this would be to use all years excluding the 2016 season as the training data set and the 2016 season as the prediction data set, or all years excluding 2016 season except for the 2016-HUTCH data as the training data set and the remaining three 2016 locations as the prediction data set.

For the VI data only rrBLUP was used for genomic prediction. The VI were used as a cofactor when predicting grain yield for that specific year-location-trial combination as follows:

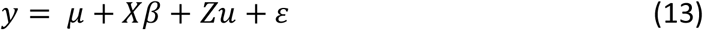

where *y* is the BLUP for grain yield, *μ* is the overall mean, *X* is a (*n x* 1) matrix of the individual observations of the VI, *β* is the fixed effects of the VI measurements, *Z* is an (*n x m*) matrix assigning markers to genotypes, *u* is a (1 *x n*) array of the random effects of the markers and *ε* is the residual error. The same cross-validation procedure as before was used.

Unless otherwise stated, all analysis took place in the R software environment using the Tidyverse suite of packages (Core Team 2020; Wickham et al. 2019) and visualised with ggplot2 (Wickham 2016). The required code can be found at: https://github.com/megzcalvert/ProgramBreeding

## Results

### Simulation

To determine the optimal parameters for the operation of the KSU winter wheat breeding program comparing current phenotypic selection methodology verse genomic prediction a simulation was created based on economic decisions. We observed that at low heritabilities of the primary trait, prediction is favored even at low prediction accuracies (Figure 1). Once the ratio between the correlated response and the expected response is greater than 1, then selection on the correlated trait is expected to give greater genetic gain than selection on the primary trait. For the KSU wheat breeding program a primary focus on genotyping is beneficial in terms of genetic gain when the narrow-sense heritability of grain yield is below 0.4, approximately 7.5 lines can be genotyped for every one line that can be phenotyped, and the prediction accuracy of the genomic prediction models is greater than 0.3.

**Figure 0.1.**
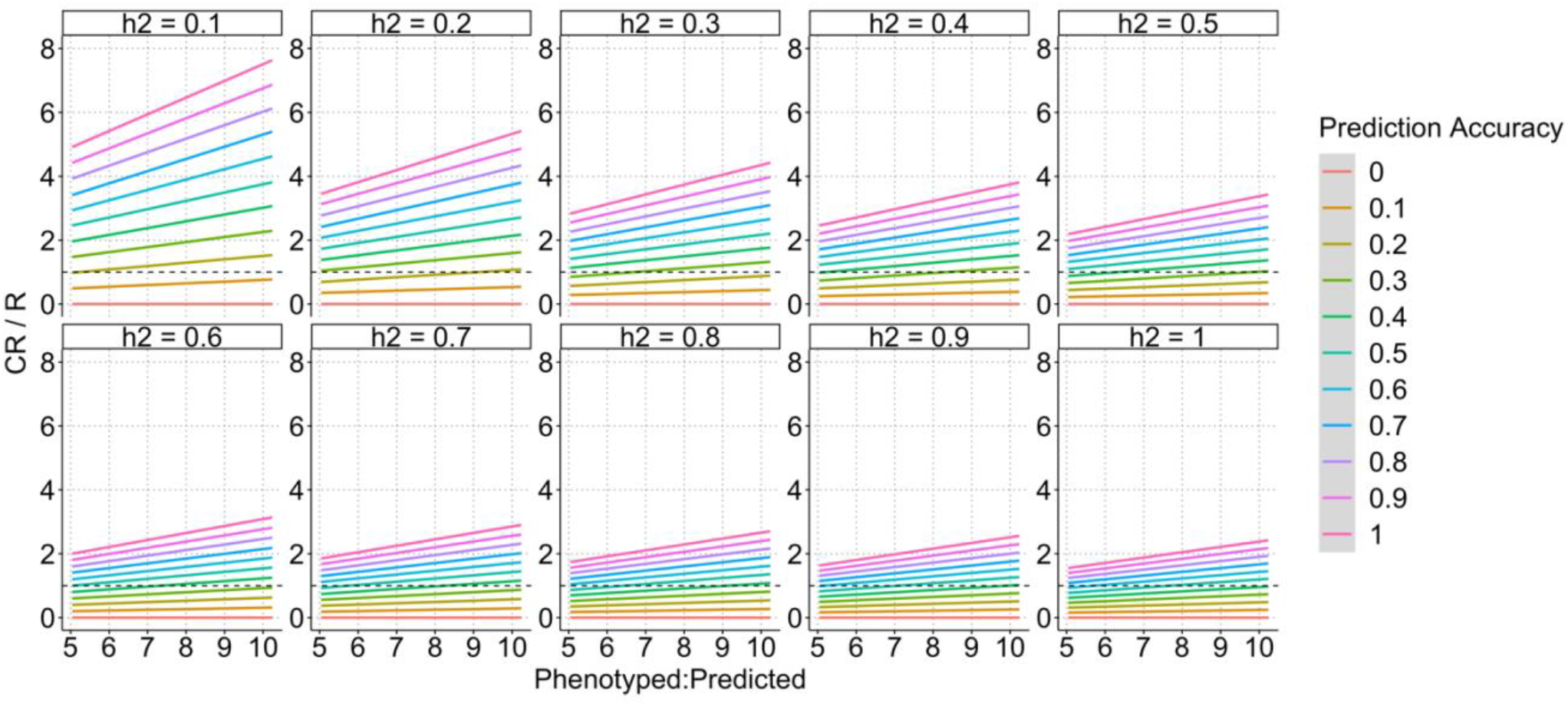
**Simulation results comparing phenotypic selection as a direct trait to selection on the genotype as a secondary trait**. **The narrow-sense heritability of the direct trait is in the panel title and the y-axis is the Correlated Response / Response of selection for the direct trait. When this ratio is above 1, represented by the dotted line, selection on the secondary trait is favored. The x-axis is the number of experimental lines that can be genotyped and predicted for every line that is phenotyped based on estimated costs. The trend of each prediction accuracy is given by the color of the line**.

### Heritability of traits

Heritability was calculated to identify the phenotypic variance that could be partitioned to the genetic component across the experimental trials. The broad sense heritabilities for each of the seasons ranged from 0.022 for the AYN to 0.441 for the PYN in the 2016 season (Figure 2). The broad-sense heritability for the 2019 PYN cannot be calculated as only one rep was planted at one location (HUTCH).

**Figure 0.2.**
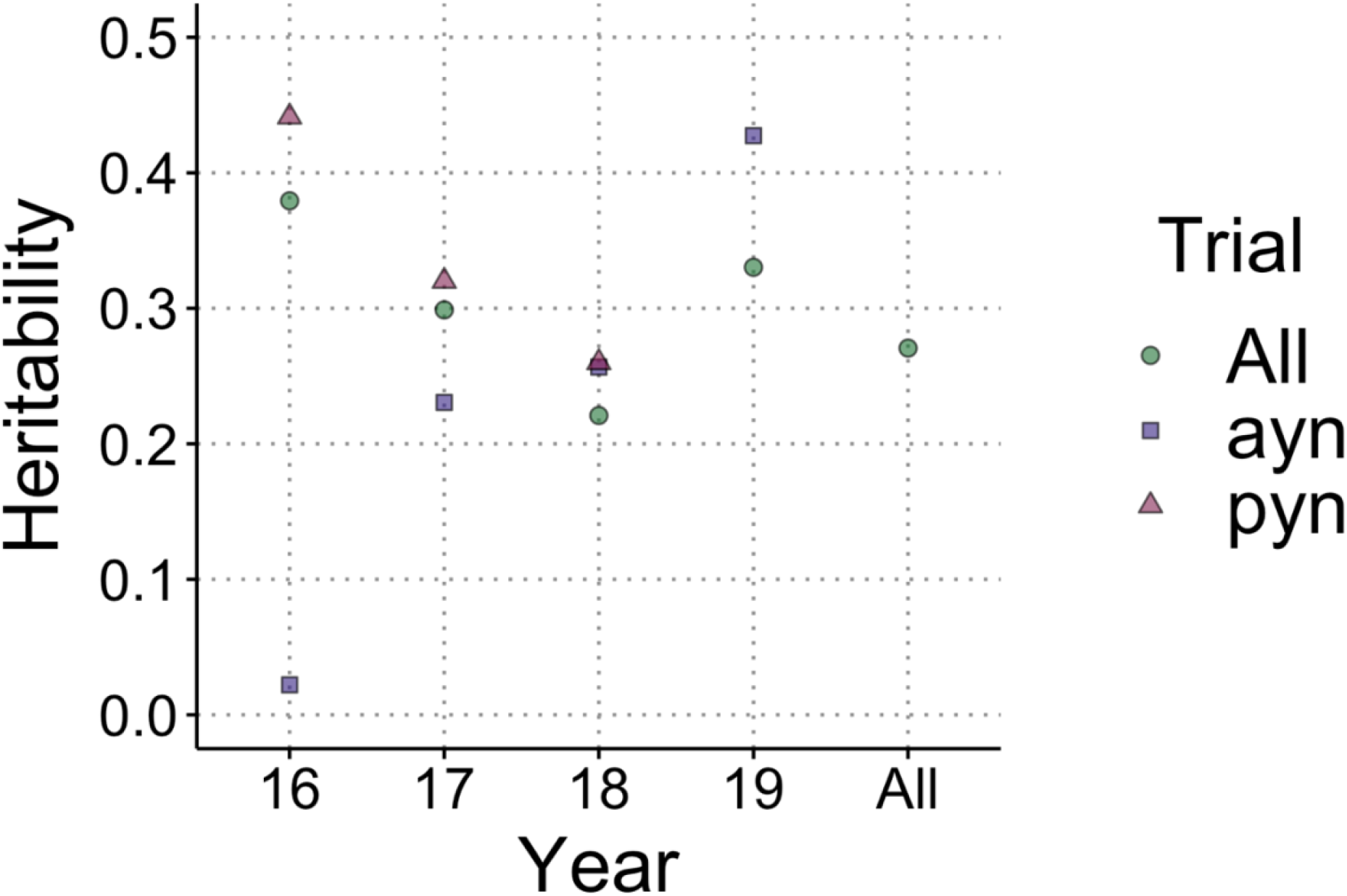
**Broad sense heritabilities for grain yield across 4 years of phenotypic data. The broad sense heritabilities are presented by season on the x-axis with each trial type denoted by the character shape. There is no PYN for the 2019 season as it was only planted in one location with one replication**.

The VIs show a moderate heritability across the seasons, with the majority being above 0.5 (Figure 3) The heritabilities are variable across the course of the season and locations but the same date-location combination shows similarity across the various VIs, indicating a large influence of ambient conditions of various days within the growing season on the accuracy of VI measurements.

**Figure 0.3.**
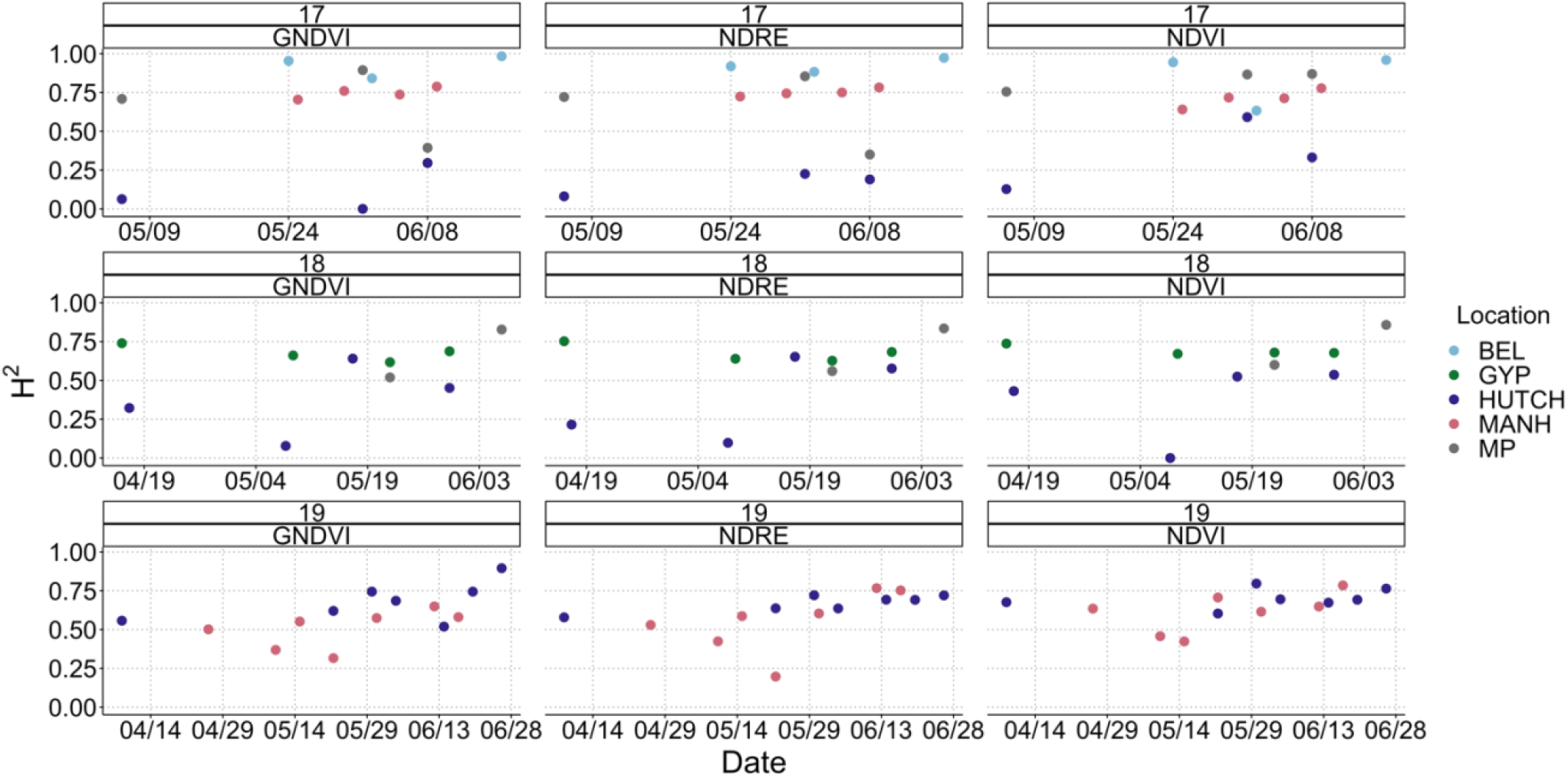
**The broad sense heritabilities for the VIs for the AYN trials planted at each location**. **The title of each plot gives the season-VI combination, with the date on which the VI was taken given by the x-axis, the y-axis is the broad sense heritability, while the color of the point determines the location. NDVI = Normalized Difference Vegetation Index, GNDVI = Green Normalized Difference Vegetation Index, and NDRE = Red Edge Normalized Difference Vegetation Index**.

The correlations between grain yield for a location-season-trial combination and the VI’s for that location-season-trial combination over the course of the season are given in Figure 4. The correlations ranged between -0.6 in the 2018 PYN at HUTCH for NDRE, to 0.69 in the 2018 AYN at MP for NDRE.

**Figure 0.4.**
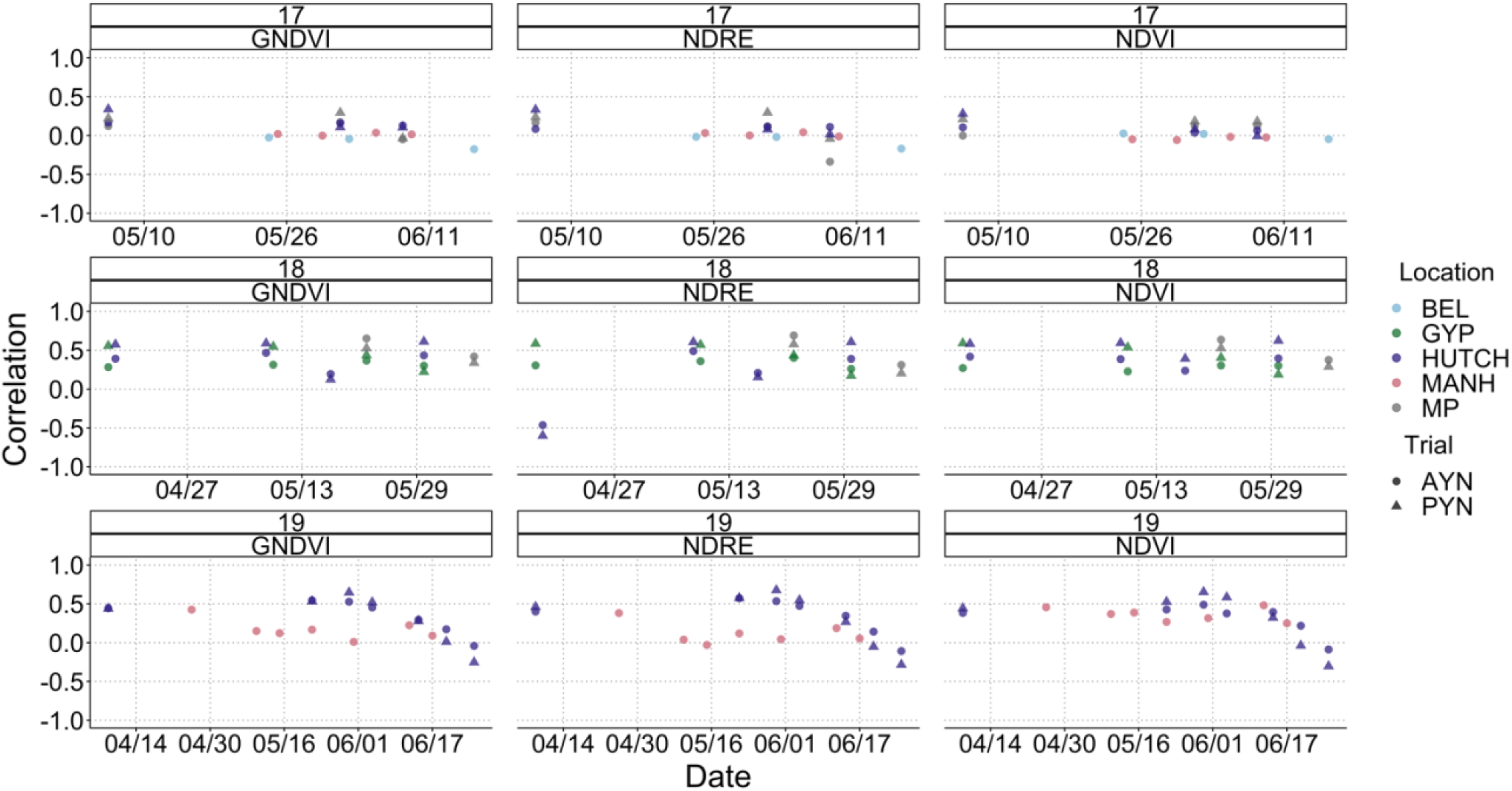
**The correlations between grain yield and VI by year-trial combinations. The title of each plot gives the season-VI combination, with the date on which the VI was taken given by the x-axis, the y-axis is the Pearson correlation coefficient, while the color of the point determines the location. NDVI = Normalized Difference Vegetation Index, GNDVI = Green Normalized Difference Vegetation Index, and NDRE = Red Edge Normalized Difference Vegetation Index**.

### GWAS

To try and determine if any of the VI had genomic regions associated with a large effect size, a GWAS was conducted. As population structure is known to have an effect on GWAS results a PCA of the 1989 experimental lines genotypic data was performed. The PCA shows no distinct population structure and the plot is bounded by the expected founder genotypes, Figure 5. There were no major QTLs for grain yield in any year, yet several of the VIs showed significant associations with regions of the genome. When these associations were further examined it was shown that they had an influence on the VI value but not GRYLD (Supplementary Figure 1). The regions that showed an association with the VI were often shared between different VI (3/4 in the 2018 season and 3/3 in the 2018 season, Supplementary Figure 1).

**Figure 0.5.**
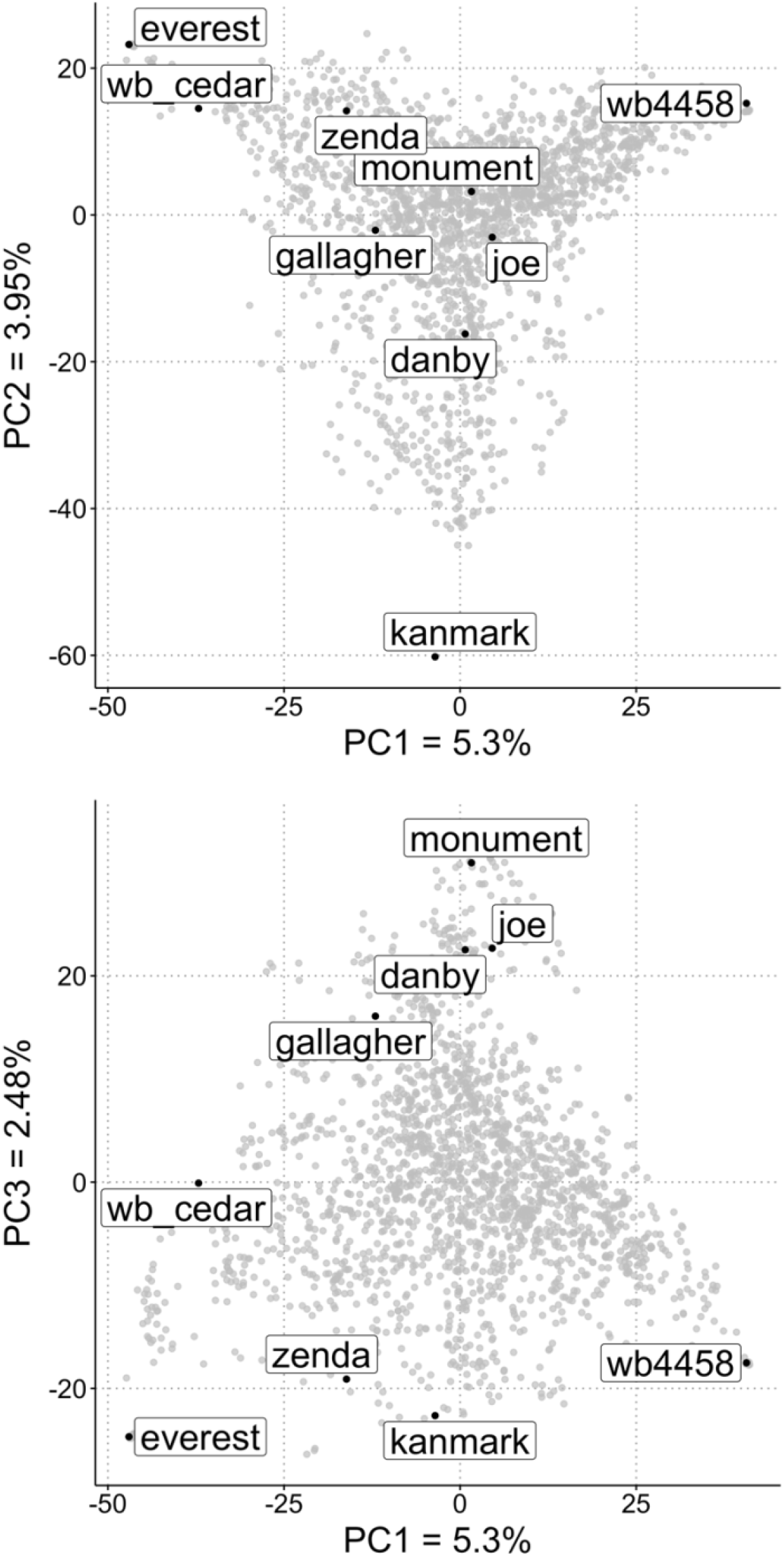
**Population structure based on genotypic information in the Kansas State winter wheat breeding program nurseries between the 2016-2019 seasons**. **The top panel shows the first two principal components (PC), while the bottom panel displays the 2nd and 3rd PC. The variance contributed by each PC is given next to the PC name. Several “founder” lines are highlighted**.

### Genomic Prediction

Based on the GWAS, GP was evaluated to predict the highly quantitative trait of grain yield. All of the tested genomic prediction models produced similar accuracies (Supplementary Table 2), we therefore focused on rrBLUP models for computational efficiently for further analysis which will be reported. The genomic prediction accuracies based on cross-validation range between 0.311 (SD = 0.079) for the 2018 season and 0.469 (SD = 0.105) for the 2017 season (Figure 6, Supplementary Table 2).

**Figure 0.6.**
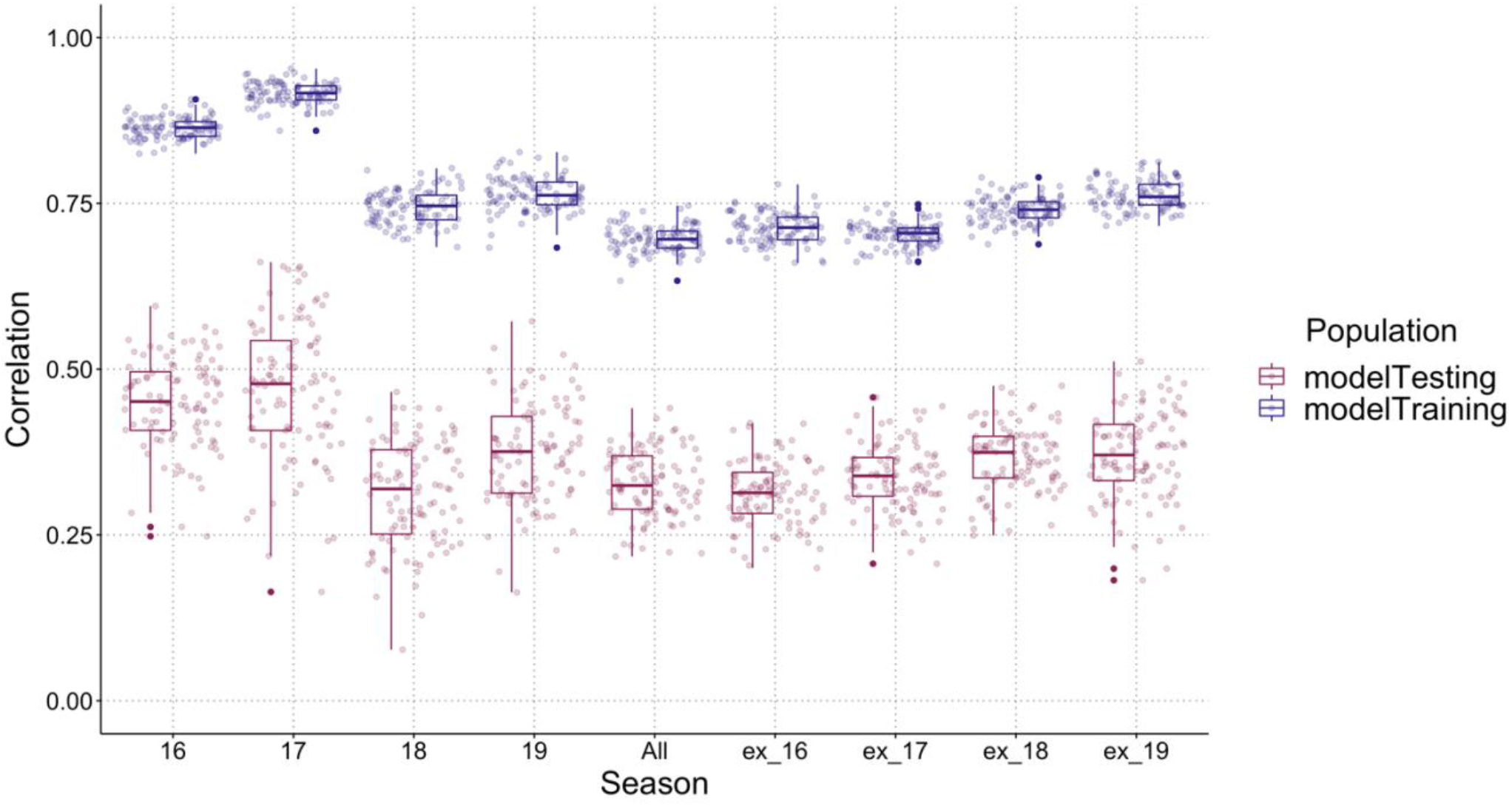
**Genomic prediction accuracies for grain yield determined by cross-validation**. **The season or seasons used in each analysis are given on the x-axis. The y-axis shows the range of possible correlations between the predicted phenotype and the observed phenotype. The color of the point determines if the individual line had been in the training or testing population of the analysis. 100 cross-validations were performed in each analysis. The modelTraining results give an indication of the model-fit whereas the modelTesting results give an indication of the predictive ability of the model. The AYN and PYN trials are included in the analysis**.

When making forward predictions the strongest correlation, -0.164, was achieved using all seasons excluding the 2017 season as the training population, and the 2017 season as the prediction population (Table 4). When predicting all locations in a single season except for HUTCH, using the data from other seasons and the data from HUTCH as the training population, the greatest accuracy achieved was 0.572 (95% CI [0.503, 0.634]) for 2019 PYN in MANH and BEL. HUTCH was chosen as the location to include in the forward predictions as it is the first location that is normally harvested in Kansas. This would be similar to harvesting one location and then using the predictions to make selections.

To evaluate a prediction approach using secondary traits, the VI were used as covariates in the prediction of grain yield. The prediction of grain yield was not improved by the addition of a covariate (Figure 7). The prediction of grain yield without a covariate was performed at the same time for comparison and was comparable to the best estimate using VI as covariate.

**Figure 0.7.**
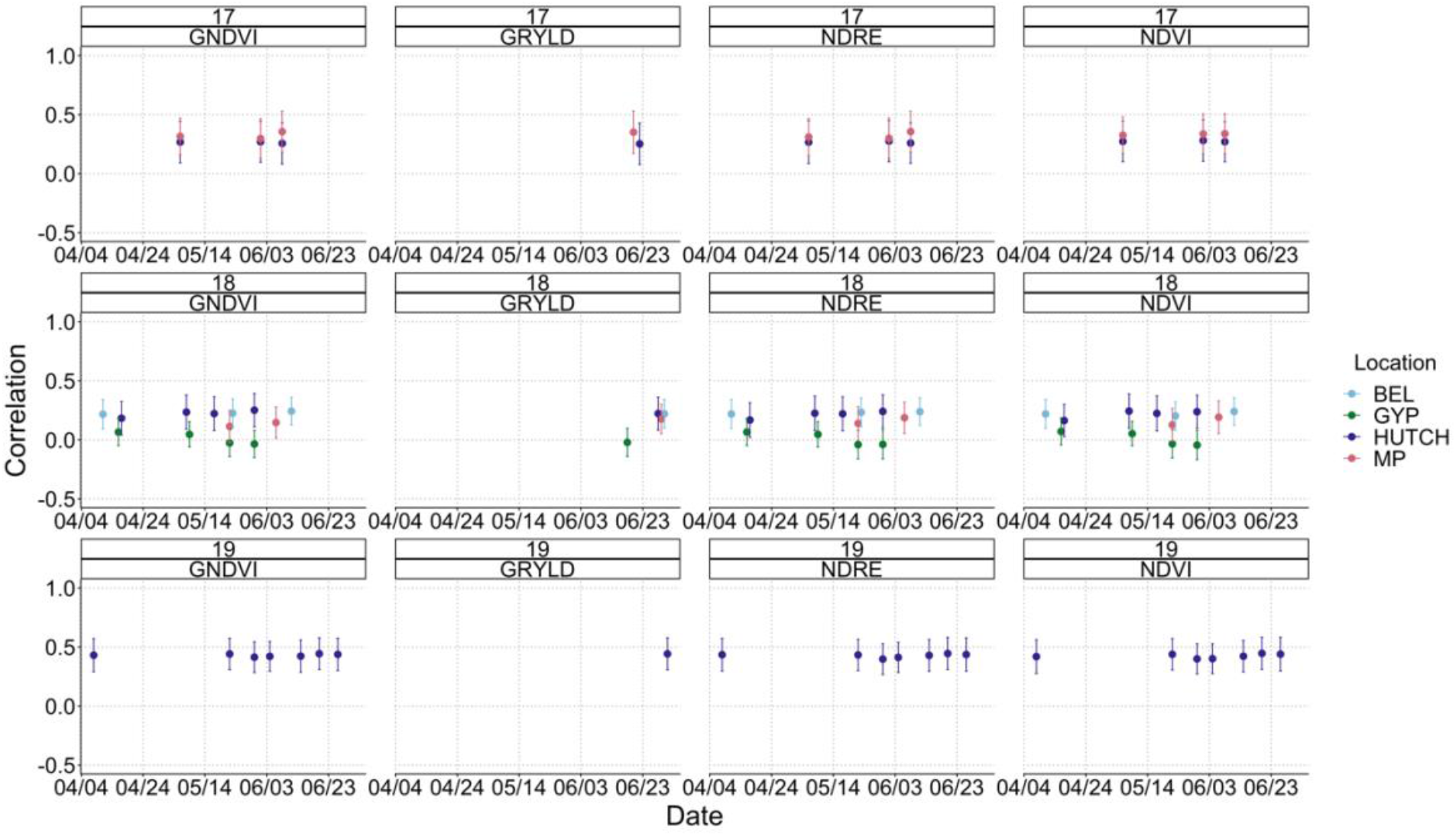
**Genomic prediction accuracies for PYN grain yield when a VI is used as a covariate determined by cross-validation**. **The season and VI are given in the strip title and the points are colored by the location. The GRYLD measurements are those for the prediction of grain yield without a covariate. The large point is the mean and the error bars give the standard deviation. NDVI = Normalized Difference Vegetation Index, GNDVI = Green Normalized Difference Vegetation Index, and NDRE = Red Edge Normalized Difference Vegetation Index**.

## Discussion

Changing global conditions require current food production systems to be resilient against unexpected events in the face of global population growth. This will require the development of crop varieties that are adapted to hotter and dryer climates. The speed at which these varieties are developed will require the adoption of new technologies that have been proven to show a positive economic investment.

This study examined the validity of using GP and other HTP techniques in the Kansas State University winter wheat breeding program based on several population and model parameters, such as heritability of the primary trait, the prediction accuracy and the selection intensity. These parameters give an indication of whether a new GP method would provide more genetic gain than traditional phenotypic selection equal resource expenditures.

### Heritability

The heritability of a trait has a significant impact on whether the trait can be selected for and the possibility for genetic gain of that trait. Traits with a lower heritability are difficult to select for regardless of which selection method is used. However, our simulation showed that traits with a low to moderate narrow-sense heritability (0-0.4) favor the use of GP. For the Kansas State University winter wheat breeding program, the overall broad-sense heritability of grain yield is below 0.3. As the narrow-sense heritability is the always less than the broad-sense heritability, the heritability favors GP under the operation cost parameters for the program.

### Prediction Accuracy

The prediction accuracy for GP when making forward predictions in the breeding program do not meet the criteria to favor GP unless very large populations are used. This is a possibility in a breeding program setting as historical grain yield data and genotypes are available for previous seasons. The experimental lines in these seasons are likely to be less related to the current lines as the parental lines in the crossing block are updated which may lower prediction accuracy, however, this is the critical assessment of prediction accuracy that is needed for implementation in the breeding program as all selections will be focused on new breeding lines into new year’s.

### HTP Prediction Accuracy

Utilizing the VI to increase the prediction accuracy of the GP models requires more optimization before it is commonly adopted. The VI need to be able to be used across locations and years for them to really have an impact on the prediction accuracy of the GP models. This will require either the manual measurement of growth stages to standardize measurements to growth stage across locations and years, or the use of another method such as thermal units to account for differences in growth stage in the training populations and prediction populations. This additional labor makes the use of VIs in GP models more costly than utilizing just the GP model. It is an additional factor that needs to be taken into account before the decision to transition to GP is made.

## Conclusion

When all parameters are considered it appears that the Kansas State University winter wheat breeding program is on the edge of a large decision. The heritability of grain yield in the breeding program as well as the cost of phenotyping compared to genotyping favor genomic prediction as the way forward for the breeding program. Yet the low accuracy of forward predictions favors the use of phenotypic selection. The forward prediction accuracy can be increased as seen in the 2019 season (Table 4), but this requires a much larger training population. With a few adjustments to the experimental design such as allowing for more replicates of the training population, and changes to the program workflow that allows for the loss of the PYN, genomic prediction could allow the KSU winter wheat breeding program to make larger genetic gains.

## Authors’ contributions

MC and BE collected data. MC and XW analyzed data. JP conceived and planned the experiments. All authors read and approved the final manuscript.

## Acknowledgements

We would like to acknowledge the undergraduate assistants, Quentin Ediger, Kyle Laessig, Jacob Ramsey, Jamie Clark, Molly Smith, Erika Kringen, Nicholas VanPelt, Bryce Teaford, who assisted with data collection. Daljit Singh, Richard Brown and Grant Williams who assisted with UAS flights and data collection.

MC was supported in part by fellowship funds from the Interdepartmental Genetics program at Kansas State University. This material is based upon work supported by the National Science Foundation under Grant No. 1238187 project “A Field-Based High-Throughput Phenotyping Platform for Plant Genetics” and the USDA NIFA International Wheat Yield Partnership grant no. 2017-67007-25933/project accession no. 1011391 project “Wheat Yield Prediction and Advanced Selection Methodologies through Field-Based High-Throughput Phenotyping with UAVs”. Any opinions, findings, and conclusions or recommendations expressed in this material are those of the author(s) and do not necessarily reflect the views of the National Science Foundation or the U.S. Dept. of Agriculture.

## Abbreviations

AYN: advanced yield nurseries
BEL: Belleville
BLUPs: Best Linear Unbiased Predictions
GEBVs: genomic estimated breeding values
GNDVI: Green Normalized Difference Vegetation Index
GP: genomic prediction
GRVI: green-red vegetation indices
GRYLD: grain yield
GYP: Gypsum
GxE: genotype x environment
HTP: high-throughput phenotyping
HUTCH: Hutchinson
KSU: Kansas State University
MANH: Manhattan
MP: McPherson
NDRE: normalized difference red edge
NDVI: normalized difference vegetation indices
PTHT: plant height
PC: principal component
PYN: preliminary yield nurseries
RKHS: reproducing kernel Hilbert spaces regression
RR: ridge regression
SNPs: Single nucleotide polymorphisms
UAS: uncrewed aerial system
VI: vegetation indices

## Supplementary Material

**Figure C.1.**
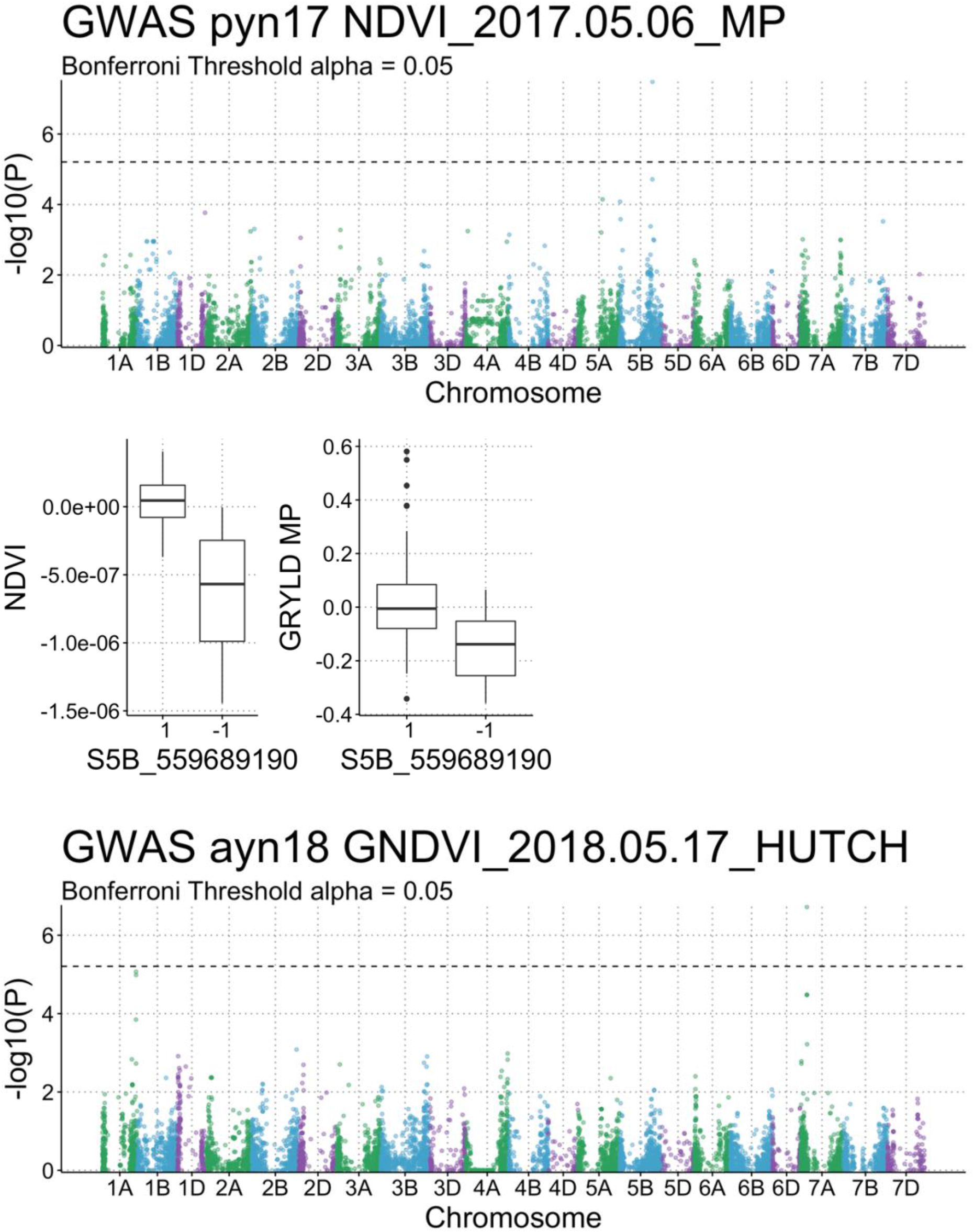
**Results of GWAS Analysis**. **The effect of the significant SNPs identified in the GWAS analysis for the specific phenotype and grain yield are given in the boxplot below the GWAS results. The points are colored by chromosome**.

**Figure C.1.**
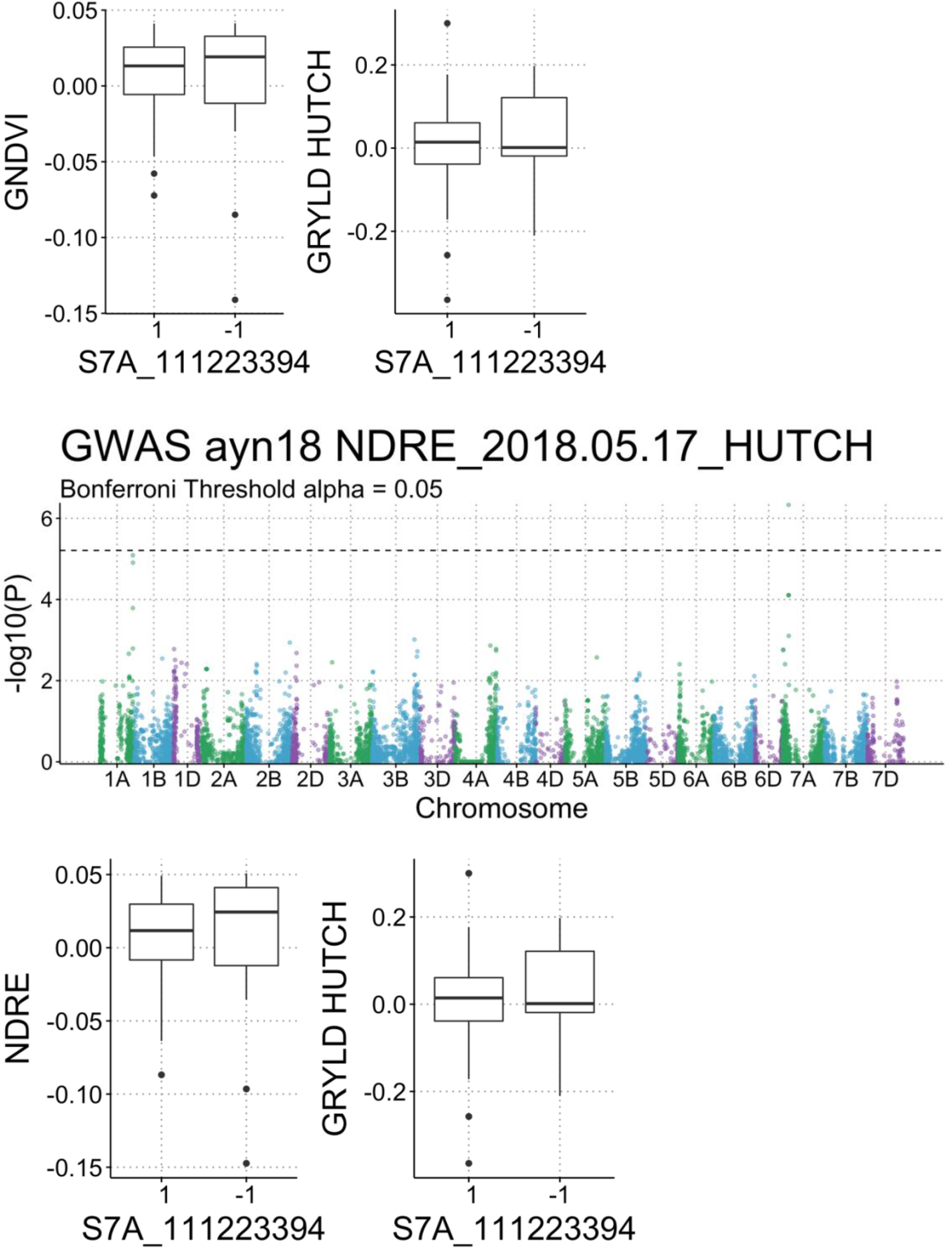
**Results of GWAS Analysis Continued**. **The effect of the significant SNPs identified in the GWAS analysis for the specific phenotype and grain yield are given in the boxplot below the GWAS results. The points are colored by chromosome**.

**Figure C.1.**
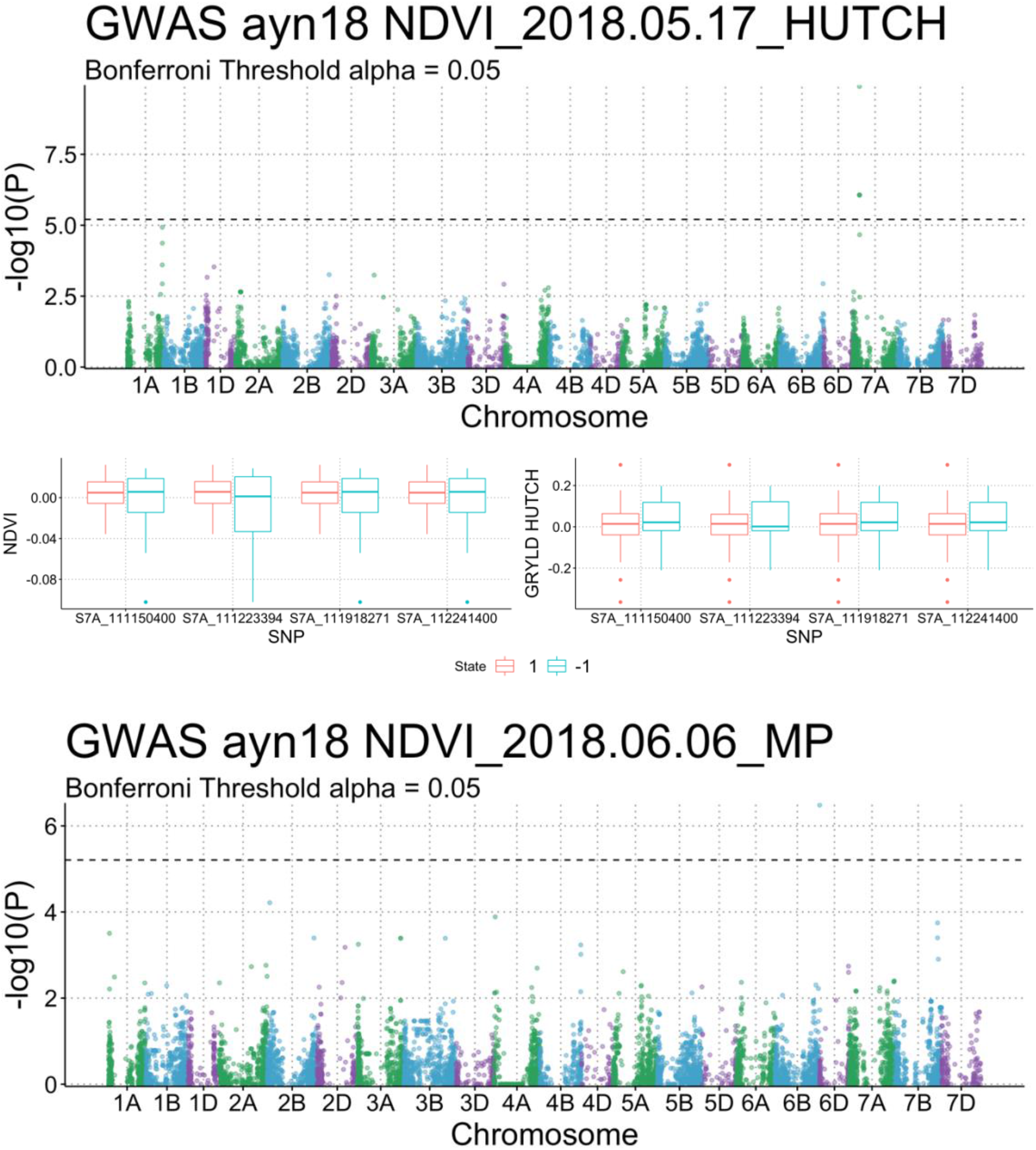
**Results of GWAS Analysis Continued**. **The effect of the significant SNPs identified in the GWAS analysis for the specific phenotype and grain yield are given in the boxplot below the GWAS results. The points are colored by chromosome**.

**Figure C.1.**
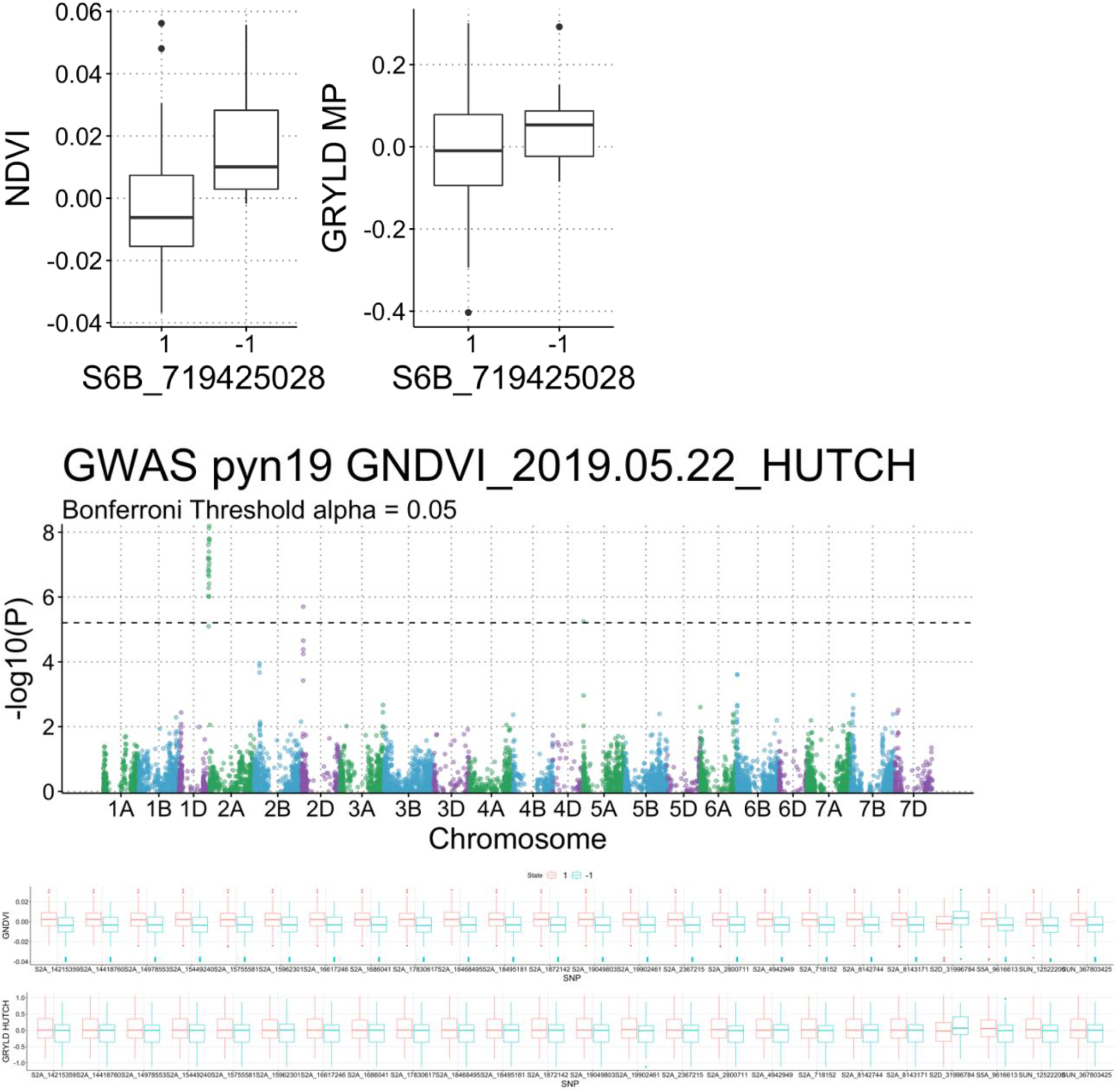
**Results of GWAS Analysis Continued**. **The effect of the significant SNPs identified in the GWAS analysis for the specific phenotype and grain yield are given in the boxplot below the GWAS results. The points are colored by chromosome**.

**Figure C.1.**
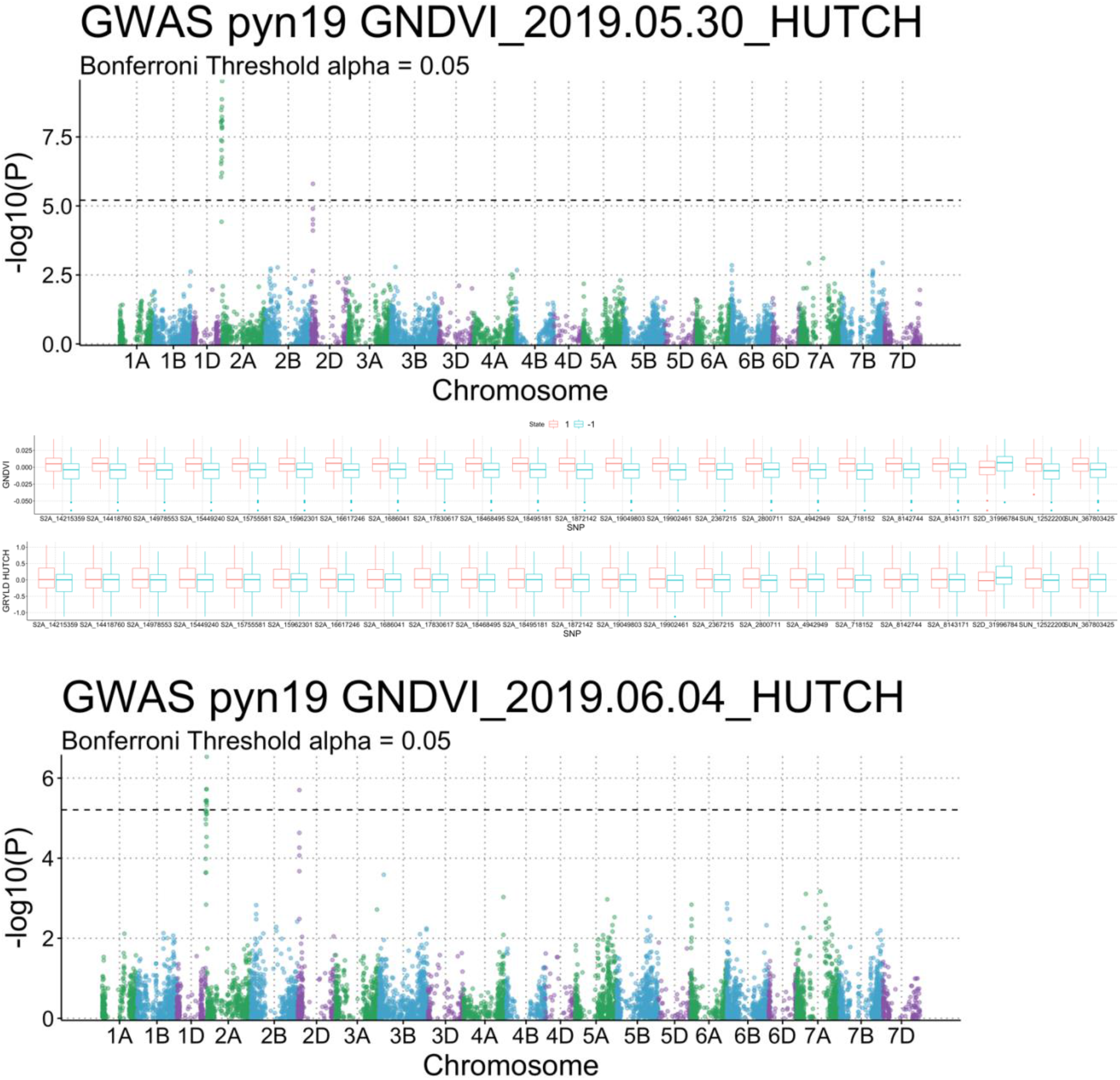
**Results of GWAS Analysis Continued**. **The effect of the significant SNPs identified in the GWAS analysis for the specific phenotype and grain yield are given in the boxplot below the GWAS results. The points are colored by chromosome**.

**Table C.1.** **Summary of simulation costs. The range over which the parameter was tested as well as the step change for each are given**.

| **Parameter** | **Range** | **Step** |
| --- | --- | --- |
| **Line Development** | \$4-40 | \$1 |
| **Genotyping Costs** | \$1-20 | \$1 |
| **Phenotyping Costs** | \$35-70 | \$5 |
| **Phenotyping Replications** | 6 | N/A |
| **Heritability of Primary Trait** | 0-1 | 0.1 |
| **Heritability of Correlated Trait** | 0.95 | N/A |
| **Prediction Accuracy** | 0-1 | 0.1 |
| **AYN Selected** | 100 | N/A |

**Table C.2.** **Accuracy of different GP models. The average correlation between the observed and GEBVs for the different genomic prediction models for 100 CV’s is given with the standard deviation in brackets**.

| **Season** | **Ridge Regression** | **Reproducing Kernel Hilbert Space** | **BayesC** |
| --- | --- | --- | --- |
| **All** | 0.3274 (0.0494) | 0.3279 (0.0492) | 0.3274 (0.0496) |
| **2015/2016** | 0.4505 (0.0700) | 0.4495 (0.0700) | 0.4500 (0.0703) |
| **2016/2017** | 0.4834 (0.1075) | 0.4843 (0.1075) | 0.4840 (0.1067) |
| **2017/2018** | 0.2963 (0.0702) | 0.2962 (0.0704) | 0.2971 (0.0702) |
| **2018/2019** | 0.3700 (0.0802) | 0.3704 (0.0793) | 0.3704 (0.0801) |
| **Excluding_2016** | 0.3197 (0.0516) | 0.3201 (0.0518) | 0.3201 (0.0526) |
| **Excluding_2017** | 0.3342 (0.0551) | 0.3345 (0.0551) | 0.3341 (0.0555) |
| **Excluding_2018** | 0.3619 (0.0533) | 0.3634 (0.0539) | 0.3623 (0.0534) |
| **Excluding_2019** | 0.3682 (0.0610) | 0.3683 (0.0611) | 0.3678 (0.0612) |

## Notes

https://github.com/megzcalvert/ProgramBreeding

### Competing Interest Statement

The authors have declared no competing interest.

### Summary of Updates

Improved figure readability Equation inclusion

